# Phenotypic and genotypic characterization of *Listeria monocytogenes* in clinical ruminant cases in Korea

**DOI:** 10.1101/2022.01.24.477645

**Authors:** Jongho Kim, Jong Wan Kim, Ha-Young Kim

**Author notes:** Address correspondence to Ha-Young Kim, or.

## Abstract

*Listeria monocytogenes* is a foodborne human and veterinary pathogen. This study aimed to determine the phenotypic and genotypic characteristics of *L. monocytogenes* isolates from clinical cases of Korean ruminants. We collected 24 *L. monocytogenes* isolates from clinical cases with caprine neurological symptoms and bovine abortion. The most prevalent serotypes were 4b (IV_b_), 1/2a (II_a_; II_c_), and 1/2b (II_b_). All isolates, including two found in humans, formed three genetically diverse pulsed-field gel electrophoresis clusters according to serotype, lineage, and sequence type. The most prevalent sequence type was ST1, followed by ST365 and ST91. *L. monocytogenes* isolates from ruminant listeriosis were resistant to oxacillin and ceftriaxone. These clinical ruminant isolates showed diverse lineage, serotype (serogroup), and sequence type characteristics. Considering that the atypical sequence types exhibited clinical manifestations and histopathological lesions, further study is needed to elucidate the pathogenicity of genetically diverse ruminant *L. monocytogenes* isolates. Furthermore, continuous monitoring of antimicrobial resistance is required to prevent the emergence of *L. monocytogenes* strains resistant to common antimicrobials.

## INTRODUCTION

Listeriosis caused by *Listeria monocytogenes* causes high morbidity and mortality primarily in ruminant farms. Listeriosis in ruminants manifests in two major forms: septicemia and (rhomb)encephalitis; the septicemic form of listeriosis can result in fetal infection and subsequent abortion (1). A Danish study showed that the diagnostic rate of *L. monocytogenes* in bovine abortion was 1.2% (2). In small ruminant surveillance studies conducted in Slovenia and Switzerland, the incidence rates of listeria rhombencephalitis and central nervous system disease in bovine adults were 5.3 and 26.3 cases per 100,000, respectively (3). In humans, *L. monocytogenes* is a serious foodborne pathogen causing various clinical conditions, including septicemia, abortion, gastroenteritis, and encephalitis (4).

Thirteen *L. monocytogenes* serotypes (1/2a, 1/2b, 1/2c, 3a, 3b, 3c, 4a, 4ab, 4b, 4c, 4d, 4e, and 7) have been identified, with lineages I (serotypes 1/2b, 4b, and 3b) and II (serotypes 1/2a, 1/2c, and 3c) more frequently isolated from ruminant listeriosis cases than lineages III or VI (1). Lineage I is most frequently isolated from human and ruminant encephalitis (1,5). Among the 13 serotypes, 1/2a and 3a have been associated with non-encephalitic infections (4). Using a multiplex PCR method to overcome the major drawbacks of conventional serotyping, these serotypes were reclassified into five serogroups: II_a_ (1/2a and 3a), II_b_ (1/2b, 3b, and 7), II_c_ (1/2c and 3c), IV_a_ (4a and 4c), and IV_b_ (4ab, 4b, 4d, and 4e) (6). Among them, PCR serogroup II_a_ is the most prevalent in both fetal infection and neurologic cases in ruminant listeriosis (1). Furthermore, *L. monocytogenes* serotypes 4b, 1/2a, and 1/2b were primarily responsible for the human cases of listeriosis in the EU in 2013 and USA between 2011 and 2016 (6,7). However, serotyping has a limited discriminatory ability when comparing bacterial genotypes (8). Therefore, molecular typing is mostly conducted using pulsed-field gel electrophoresis (PFGE) and multilocus sequence typing (MLST) to investigate the zoonotic sources and genetic diversity of *L. monocytogenes* isolates (1,4).

Several virulence genes responsible for bacterial pathogenesis in host cells have been identified in *L. monocytogenes* isolates (9). Among them, *inlA, inlC,* and *inlJ*, which encode internalin-like proteins, are considered to be important for the initial stage of infection; *llsX* (listeriolysin S expression), *plcA* (phosphatidylinositol-phospholipase C), *plcB* (phosphatidylcholine-phospholipase C), *hly* (listeriolysin O)*, lmo2672* (transcriptional regulator), and *prfA* (transcriptional regulator) are crucial for the development of human listeriosis (9,10).

Listeriosis is susceptible to most antimicrobials used to treat gram-positive bacteria (11,12). A combination of β-lactam antimicrobials, such as penicillin or ampicillin, and aminoglycosides, such as gentamicin, has been used as the first choice for human listeriosis. Trimethoprim-sulfamethoxazole, tetracycline, and erythromycin have been used as alternative therapies for listeriosis (9,11,13). However, several *L. monocytogenes* isolates are resistant to these antimicrobials (9,10,13,14). In Korea, *L. monocytogenes* isolates from swine and bovine carcasses have showed resistance to the recommended antimicrobials, such as penicillin (93.3%), ampicillin (26.7%), and tetracycline (20.0%) (15). There are currently no studies on the antimicrobial resistance of *L. monocytogenes* isolates from clinical ruminant cases.

The aim of the present study was to determine the phenotypic and genotypic characteristics of *L. monocytogenes* isolates from clinical cases of Korean ruminants. Furthermore, we monitored antimicrobial resistance and determined empirical antimicrobials of *L. monocytogenes* isolates from clinical cases of ruminants.

## MATERIALS AND METHODS

### Isolates and DNA extraction

For differential diagnosis, 360 stillborn bovine fetuses (2015–2019) and 99 goats (2013– 2019) presenting with listeriosis-related symptoms, including neurological signs, abortion, and sudden death, were requested from farm owners. All isolates were subjected to biochemical identification using the VITEK^®^ II system (BioMérieux, Marcy l’Etoile, France) and VITEK^®^ MS system (matrix-assisted laser desorption ionization time-of-flight mass spectrometry; MALDI-TOF MS, BioMérieux). The isolates were confirmed by the amplification of *hly* (16) and used to inoculate brain-heart infusion broth (Becton Dickinson, Sparks, MD, USA). These solutions were then aerobically cultured for 18 h at 37 °C. Genomic DNA was extracted using the Maxwell^®^ RSC instrument (Promega, Madison, WI, USA) with the Maxwell^®^ RSC Blood DNA kit (Promega) according to the manufacturer’s instructions. In addition, two human *L. monocytogenes* isolates (NCCP 14714, NCCP 15743) were obtained from the National Culture Collection for Pathogens (NCCP) of the Korea Disease Control and Prevention Agency (KDCA).

#### PCR serogrouping and conventional serotyping

Multiplex PCR was used to determine *L. monocytogenes* molecular serogroups as previously described (1). For agglutination analysis, a commercially available serotyping kit (Denka Seiken Co., Tokyo, Japan) was used to identify O and H antigens according to the manufacturer’s instructions.

### Detection of virulence genes

The 11 virulence genes *inlA*, *inlC*, *inlJ*, *lmo2672*, *llsX*, *prfA*, *plcA*, *hlyA*, *mplA*, *actA*, and *plcB* were identified from *L. monocytogenes* isolates using PCR, as described previously (9).

### PFGE

PFGE of the *L. monocytogenes* isolates was performed using a previously described protocol, with some modifications (15). Briefly, bacterial DNA was digested using the restriction enzyme *AscI* (Thermo Scientific, Waltham, MA, USA) for at least 5 h at 37°C. Size separation of restricted DNA fragments was performed using a CHEF MAPPER XA system (Bio-Rad, Richmond, CA, USA) with 1% SeaKem gold agarose gel for 21 h. *XbaI*-digested *Salmonella enterica* serotype Braenderup H9812 DNA was used as a standard to compare each band of the *L. monocytogenes* isolates. The banding patterns were then interpreted and compared using GelCompar II software (Applied Maths, Sint-Martens-Latem, Belgium). A combined dendrogram was analyzed using the unweighted pair group method with arithmetic mean with Dice correlation coefficient and 1% tolerance.

### MLST

*Listeria monocytogenes* isolates were subtyped using the MLST scheme, as described in the Institut Pasteur MLST database (http://bigsdb.pasteur.fr/listeria). Sequence data for seven housekeeping genes (*abcZ*, *bglA*, *cat*, *dapE*, *dat*, *ldh*, and *lhkA*) were analyzed using an ABI Prizm 3730XL analyzer (Applied Biosystems, Foster City, CA, USA) at Macrogen (Macrogen Inc., Seoul, Korea). Allelic profiles and phylogenetic lineage corresponding sequence types (STs) with clonal complexes (CC) were determined according to the criteria in the MLST Pasteur database. MEGA 7.0 (Pennsylvania State University, State College, Pennsylvania, USA) was used to construct a neighbor-joining phylogenetic tree from the concatenated alignment of allele sequences, with Kimura’s two-parameter model and 1,000 replicates (17). Concatenated sequences obtained from clinical ruminant isolates in this study and human *L. monocytogenes* strains from the NCCP of KDCA were phylogenetically analyzed. ST72 from lineage III was chosen as the root. For comparison with common clinical ruminant *L. monocytogenes*, ruminant listeriosis-associated STs from Europe and common human listeriosis-associated STs, such as ST2, ST6, and ST37, were used for phylogenetic analysis.

### Antimicrobial susceptibility test

Antimicrobial susceptibility testing of the *L. monocytogenes* isolates was performed using a Sensititre GPN3F plate (Trek Diagnostic Systems, Cleveland, USA), which contained 18 antimicrobials, following the manufacturer’s instructions. The antimicrobials and their ranges in the GPN3F plate are shown in Table S1. The antimicrobial resistance of the isolates was determined according to the guidelines of the Clinical and Laboratory Standards Institute, adopting the criteria for *L. monocytogenes* and other gram-positive bacteria, except for ampicillin, erythromycin, and penicillin, for which the European Committee on Antimicrobial Susceptibility Testing breakpoints were used (18,19).

### Statistical analysis

Statistical testing was performed using GraphPad Prism (v5.01; GraphPad Software, San Diego, CA, USA). A 2 × 2 contingency table using Fisher’s exact test was used to assess associations among serotypes and virulence genes. Statistical significance was set at *P* < 0.05.

## RESULTS

### Clinical features of caprine and bovine listeriosis

The prevalence rates of listeriosis in bovine and caprine samples were 2.5% (9/360) and 16.2% (16/99), respectively. Among them, 24 *L. monocytogenes* isolates were examined in this study, including 8 bovine fetal abortion cases and 16 caprine listeriosis. According to the clinical features of the ruminant listeriosis cases observed (Table 1), most cases (66.7%, 16/24) occurred in the spring between March and May, and all bovine abortions caused by *L. monocytogenes* occurred during late-term pregnancy (8–9 months). *L. monocytogenes* was the only pathogen in all caprine listeriosis cases that presented neurological symptoms, including circling. Furthermore, all caprine cases showed suppurative meningoencephalitis with microabscesses in the brainstem. In bovine abortion cases, *L. monocytogenes* was also the only pathogen identified, with the exception of two cases that displayed co-infection with *Leptospira* species and *Neospora caninum*, respectively. Among the eight bovine abortion cases, suppurative placentitis was confirmed in two cases, and histopathological lesions were not identified in other cases.

**Table 1.**
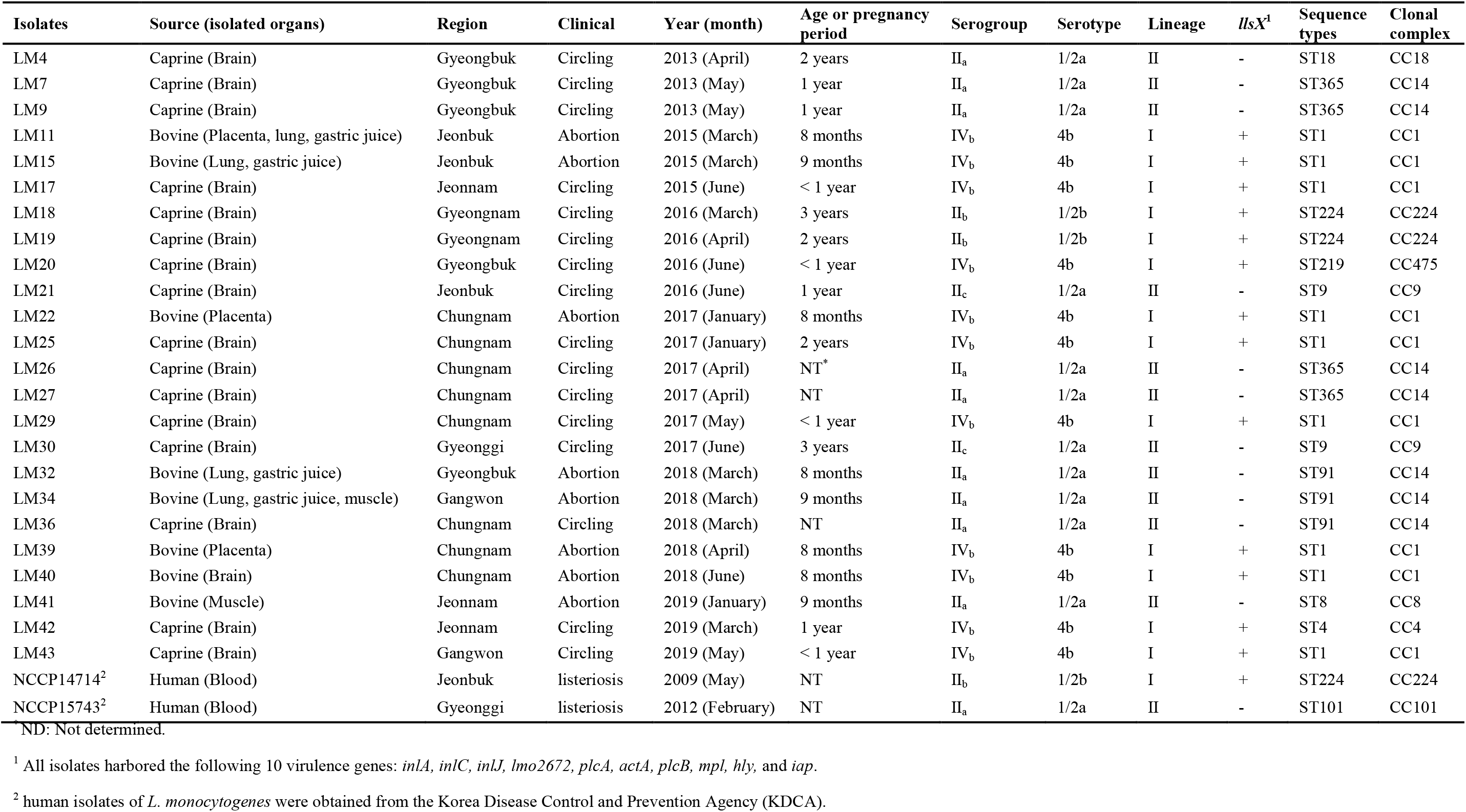
Clinical features and characterization of ruminant (n = 24) and human (n=2) *Listeria monocytogenes* isolates in Korea

### PCR serogroups and serotypes of *L. monocytogenes* isolates

Among the 24 *L. monocytogenes* isolates, PCR serogroup IV_b_ (n = 11, 45.8%) was the most prevalent, followed by II_a_ (n = 9, 37.5%), II_b_ (n = 2, 8.3%), and II_c_ (n = 2, 8.3%), with most belonging to 1/2a (n = 11, 45.8%) and 4b (n = 11, 45.8%) serotypes. The remaining two isolates were 1/2b (Table 1). Interestingly, serotypes 4b and 1/2b corresponded to serogroups IV_b_ and II_b_, respectively. However, serotype 1/2a included the two serogroups, II_a_ and II_c_.

### Virulence genes expressed in *L. monocytogenes* isolates

All isolates expressed the virulence genes *inlA*, *inlC*, *inlJ*, *lmo2672*, *plcA*, *actA*, *plcB*, *mpl*, *hly*, and *iap* (Table 1). In contrast, *llsX-*encoding listeriolysin was expressed in 13 of the evaluated isolates (54.2%); however, *llsX* was not observed in isolates with the serotype 1/2a and was significantly correlated with serotype 4b (*P* < 0.001).

### PFGE analysis of *L. monocytogenes* isolates

Among the 24 *L. monocytogenes* isolates, 14 PFGE banding patterns were obtained, and the isolates were grouped into three clusters according to lineage, serotype, and ST, regardless of geographical and host source (Fig. 1). Cluster I had a similarity level of 76.6%, which included six *L. monocytogenes* isolates (lineage I, serotype 4b, and ST1). Cluster III (n = 4) included two isolates (lineage I, serotype 1/2b, and ST224) with 92.9% similarity and two other isolates (lineage I, serotype 4b, and ST1) with 96.6% similarity. All the *L. monocytogenes* isolates in Cluster II belonged to lineage II and serotype 1/2a. Cluster II (similarity level of 56.0%) contained three ST91 and two ST9 isolates.

**Figure 1.**
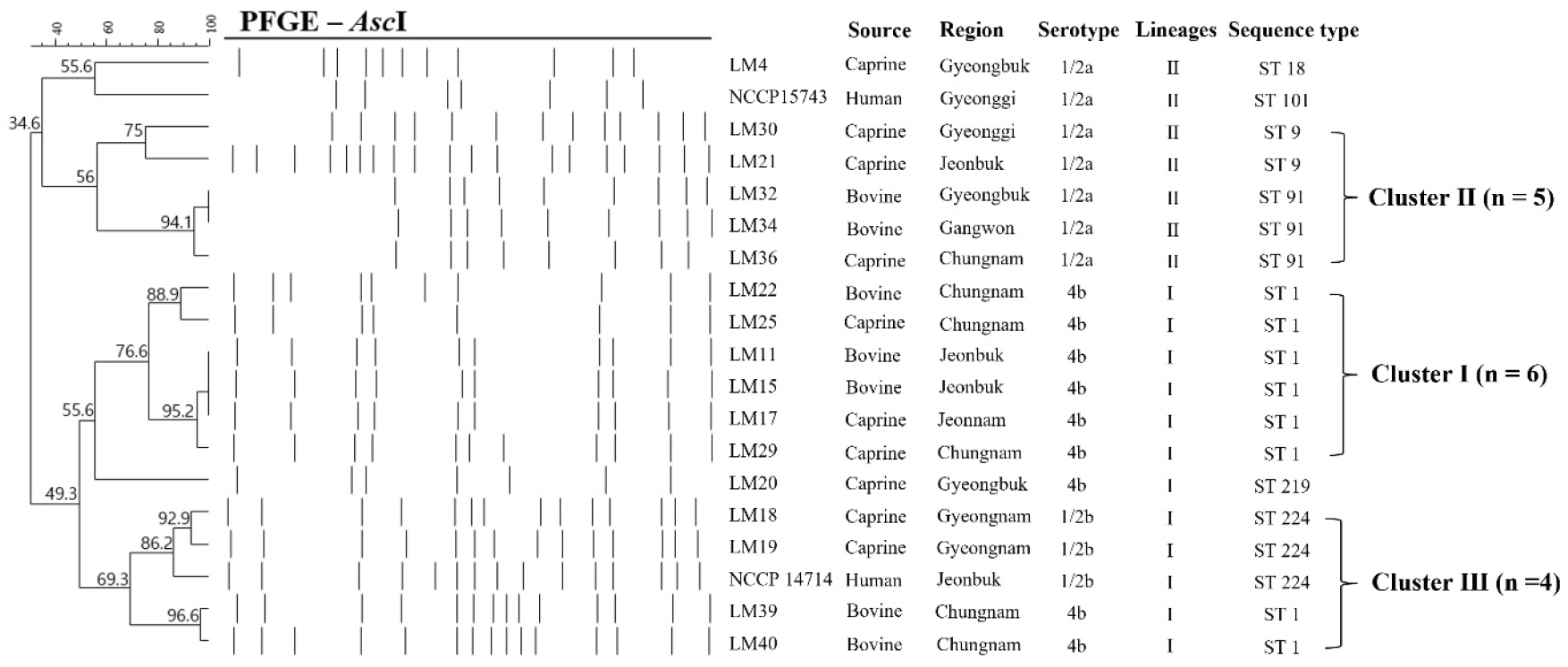
PFGE dendrogram generated using the unweighted pair group method with the arithmetic mean method from *Asc*I of *L. monocytogenes* isolates from ruminant listeriosis cases.

### MLST of L. monocytogenes isolates

Among the 24 ruminant *L. monocytogenes* isolates, the most common ST (CC) was ST1 (CC1; 37.5%, n = 9/24), followed by ST365 (CC14; 16.7%, n = 4/24) and ST91 (CC14; 12.5%, n = 3/24) (Fig. 2). The most prevalent lineage in ruminant isolates was lineage I (66.7%, n = 13), which included ST1, ST4, ST219, and ST224. The remaining STs included ST365, ST91, ST9, ST8, and ST18 and belonged to lineage II (33.3%, n = 11) (Fig. 2). The distribution of STs and genetic relatedness of *L. monocytogenes* isolates, including a clinically common bovine ST in France and Slovenia (ST37) and human isolates from common STs (ST2 and ST6), are illustrated in the phylogenetic tree (Fig. 3). STs from lineage I were more closely genetically related than those in lineage II (Fig. 3).

**Figure 2.**
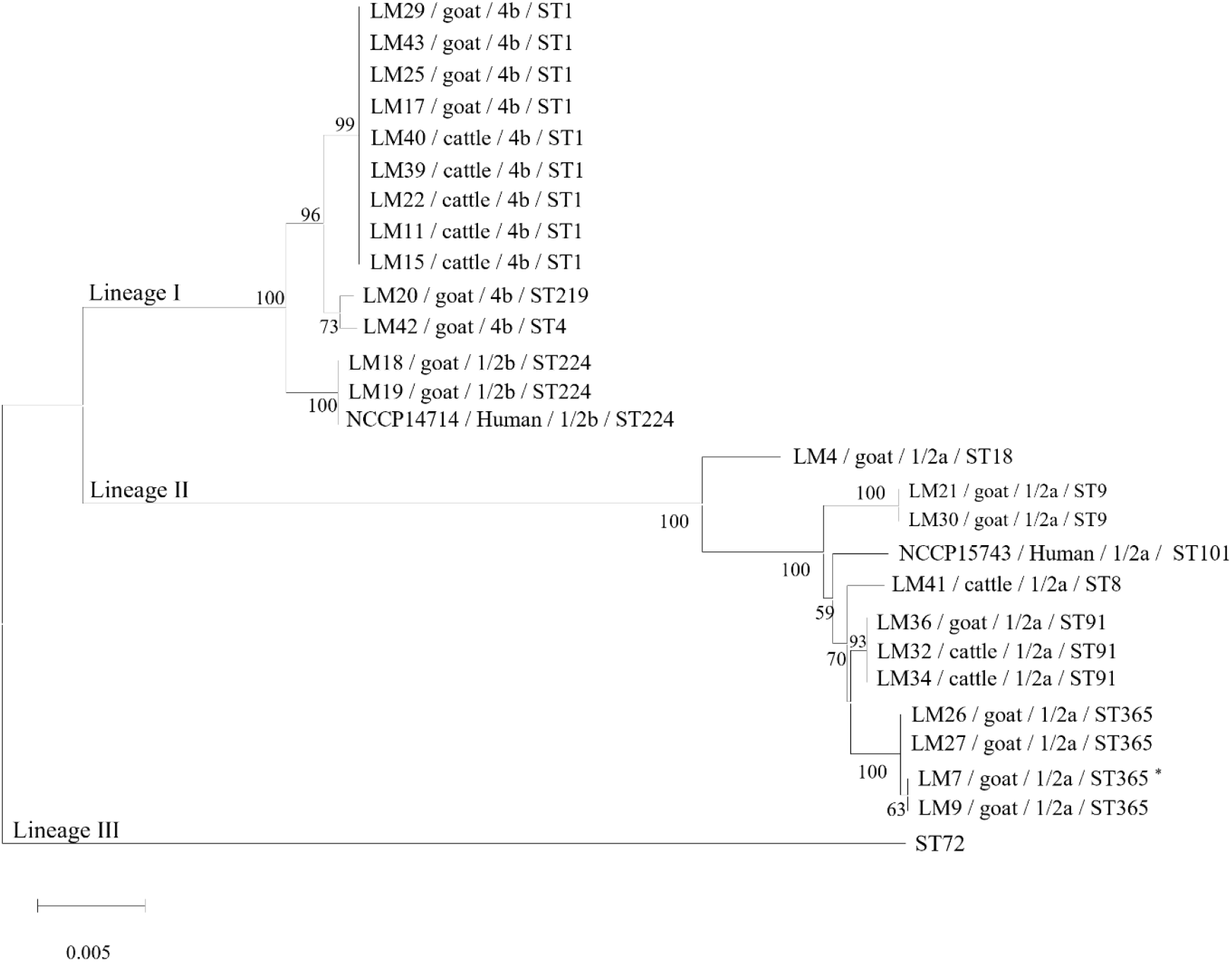
Phylogenetic analysis of concatenated MLST loci using the neighbor-joining method with 1,000 replicates. Strain names, hosts, serotypes, and sequence types are indicated. * One base difference was found (^9^T → ^9^C). The closest allele was 3. ST, sequence type.

**Figure 3.**
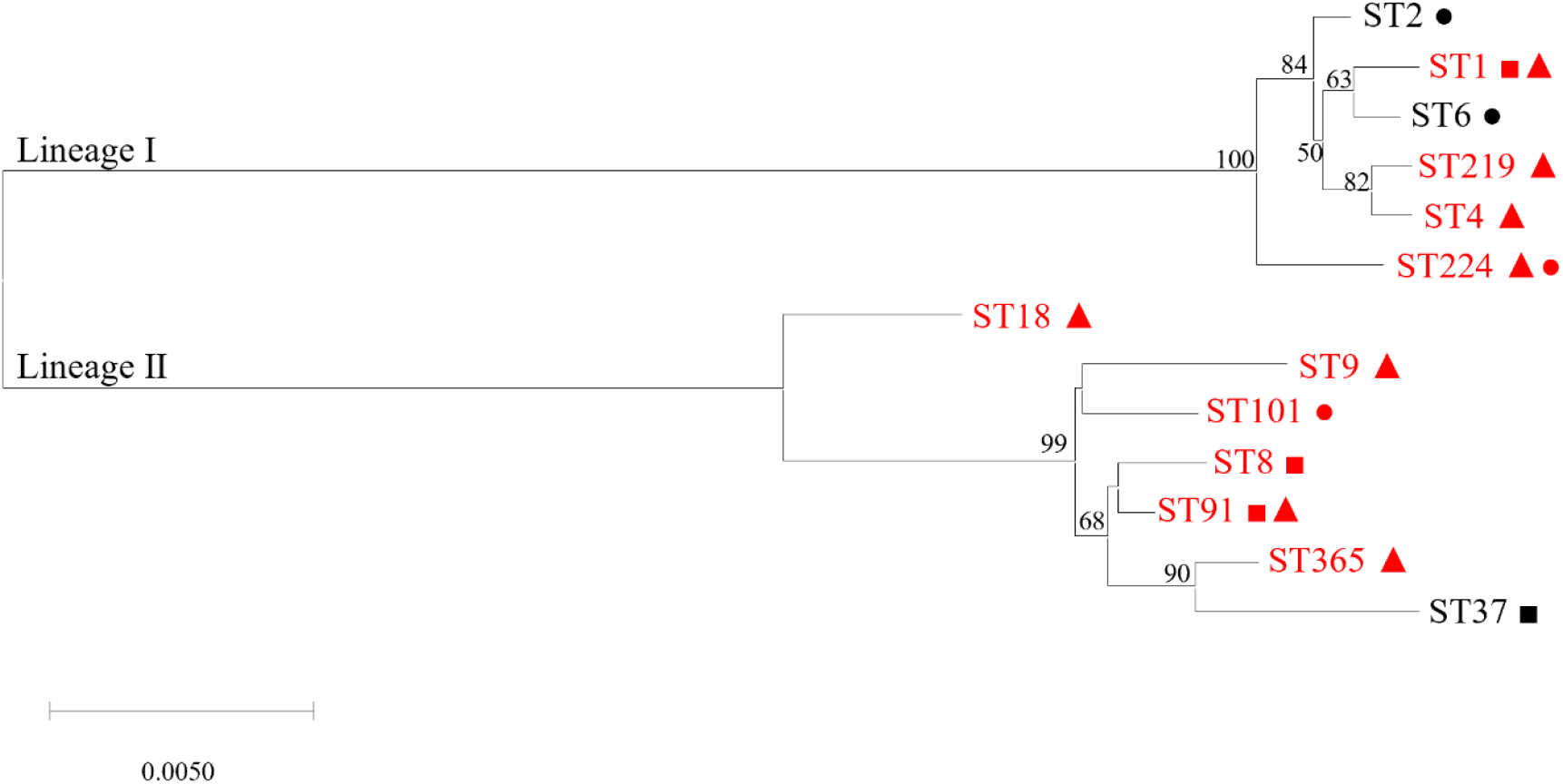
Phylogenetic analysis of concatenated MLST loci using the neighbor-joining method with 1,000 replicates. Sequence types (STs) represented in the phylogeny tree were identified from this study and obtained from the MLST database as references. STs identified in this study are represented in red. STs isolated from bovine, caprine, and human listeriosis cases are indicated with a square (▪), triangle (▴), and circle (•), respectively. ST, sequence type.

### Antimicrobial resistance in ruminant *L. monocytogenes* isolates

The antimicrobial resistance patterns of the *L. monocytogenes* isolates are shown in Table 2. All isolates tested were susceptible to 11 antimicrobials; however, many isolates were resistant to oxacillin (70.8%) and ceftriaxone (62.5%). Furthermore, several isolates showed intermediate resistance to clindamycin (58.3%), ceftriaxone (29.2%), ciprofloxacin (29.1%), and linezolid (12.5%). We did not identify isolates exhibiting multiresistance to the antimicrobials tested.

**Table 2.**
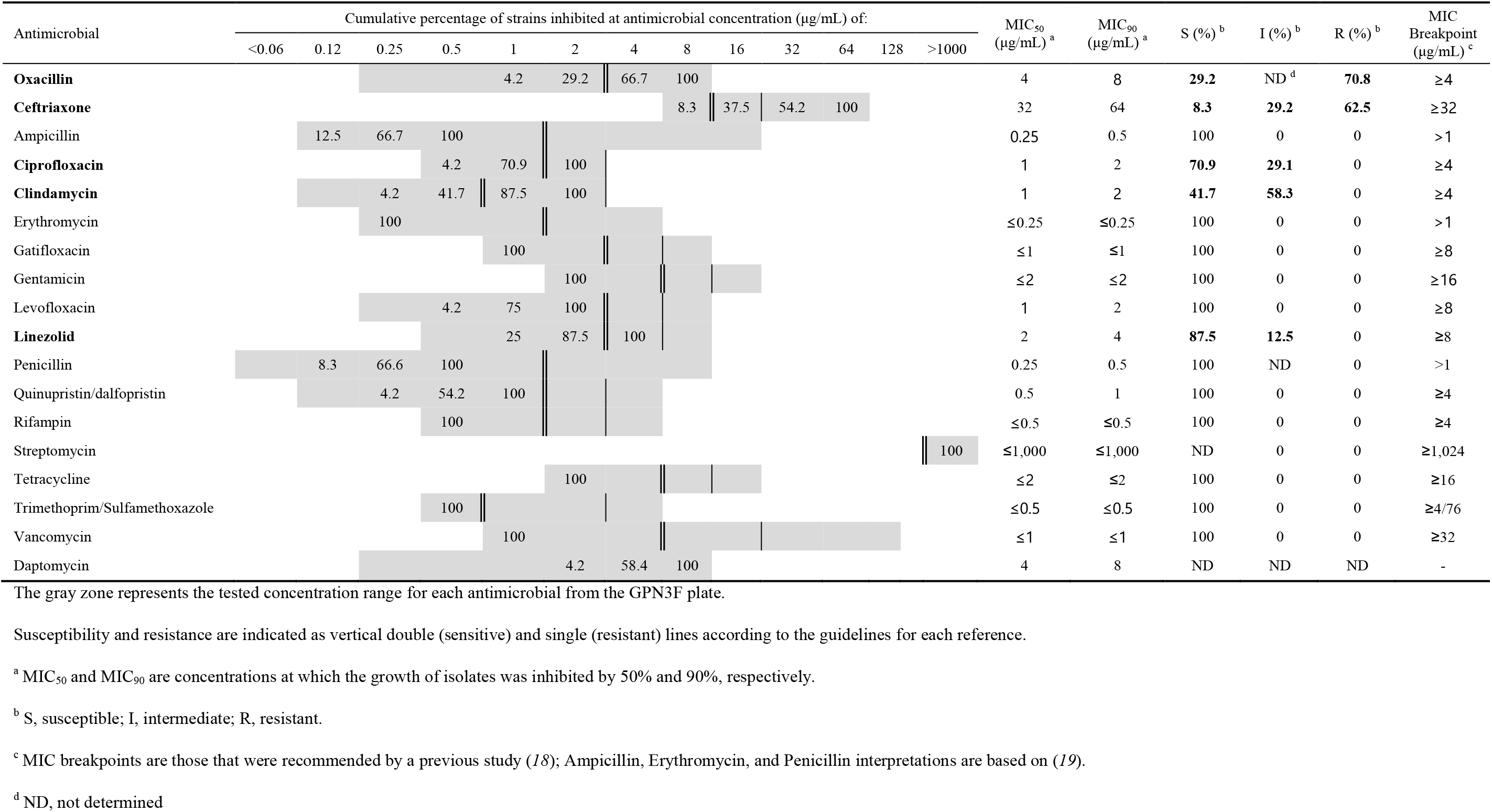
Antimicrobial susceptibility and cumulative percentage of *L. monocytogenes* (n = 24) inhibited by 18 antimicrobials

## DISCUSSION

*L. monocytogenes* is a major foodborne pathogen associated with high morbidity and mortality in infected animals. Although there have been several studies on *L. monocytogenes* isolates from foods and environments, characterization studies of ruminant *L. monocytogenes* isolates are limited in Korea (15,20). Moreover, there is little information available on the prevalence, serotypes, antimicrobial resistance, and molecular characteristics of *L. monocytogenes* isolates from ruminant clinical cases worldwide. In our study, the diagnostic rate of bovine fetal abortion was slightly higher than that in a Danish study (2). The high incidence rate in goat encephalitis was hypothesized to be from the sample collection of listeriosis cases that presented neurological symptoms. In a previous study, PCR serogroup II_a_ (serotypes 1/2a and 3a) was the dominant serogroup from ruminant listeriosis cases in the USA (1). Conversely, PCR serogroup IV_b_ and serotype 4b, which are responsible for human listeriosis, were also the most prevalent type identified from ruminant clinical listeriosis cases analyzed in this study (Table S2). The identified serotypes were limited to serotypes 1/2a, 1/2b, and 4b, which are the predominant serotypes that cause human listeria infection (21,22).

All the *L. monocytogenes* isolates examined in this study expressed 10 virulence genes, indicating that these isolates may be potentially pathogenic to humans. Indeed, *llsX* is important for the pathogenesis of human listeriosis and was expressed in the isolates that belonged to serotype 4b (*P* < 0.001) and 1/2b (10). Therefore, the serotype 4b strains from ruminant listeriosis could present a potential risk factor for human listeriosis.

Our PFGE results revealed that *L. monocytogenes* isolates from ruminant listeriosis cases were correlated with serotype and ST rather than host source or geographical origin. Based on these results, three clusters that demonstrated 55% similarity showed high genetic variation. Interestingly, two isolates (serotype 1/2b/ST224) from caprine listeriosis had relatively high similarity (85.2%) with the human blood strain NCCP 14714, which belongs to the same serotype (PCR serogroup), virulence profile, and ST reported in two caprine isolates. These caprine and human listeriosis isolates were not related to the outbreak year or geographical origin. Further epidemiological investigations and more powerful molecular characterization methods, such as whole-genome sequencing, are required to elucidate the possibility of zoonotic listeriosis and genetic relatedness of these cases.

We found that *L. monocytogenes* lineage I (n = 13, 54.2%) was more frequently detected than lineage II (n = 11, 45.8%), which was consistent with previous MLST studies of ruminant listeriosis in Europe (3,5). In contrast, a USA survey showed that lineage II was the most prevalent in ruminant listeriosis, and the number of STs was more diverse than that observed in this study (1). Moreover, ST1, ST4, and ST412 accounted for 51% of the cases, with ST1 (33.3%) being the most prevalent type in ruminant listeriosis cases in Europe (5). In another ruminant clinical case from France and Slovenia, the three most prevalent clones were CC1 (39.1%), CC4-CC217 (12.9%), and CC37 (6.0%) (3). In the USA, ST7 (15.2%) was the most frequently isolated from ruminant listeriosis, followed by ST91 (10.9%) (1). When comparing these and our results, we estimated that ST1 and ST91 were common in ruminant listeriosis despite their regional differences. Interestingly, ST365, which was the second most frequently isolated ST in this study, is mainly isolated from vegetation and rarely from ruminant listeriosis, according to the Institut Pasteur MLST database (http://bigsdb.pasteur.fr/listeria). In the phylogenetic tree, ST365 was clustered with ST37, the third most prevalent ruminant listeriosis ST in France and Slovenia cases (Fig 3). Considering the clinical manifestations and genetic relatedness with ST37, these results suggest that ST365 is a significant factor in ruminant listeriosis that should be considered in future investigations.

The lineage I clones CC1, CC4, CC2, and CC6 were described as clinically associated and hypervirulent in humans (3). However, the possibility of zoonotic infection should be noted because ST1 and ST4 strains comprised 41.7% of the cases we characterized. Moreover, according to the Pasteur MLST *L. monocytogenes* database, other STs, including ST8, ST9, ST18, ST91, ST224, ST219, and ST365 that have mainly been isolated from food and the environment have also been isolated from human infections worldwide (http://bigsdb.pasteur.fr/listeria/). Therefore, these STs may pose a risk to public health.

Although antimicrobial resistance in *L. monocytogenes* isolates has not yet been unnoticed, it is important to evaluate the effectiveness of antimicrobials and monitor the emergence of resistant strains. Fortunately, we found no multi-drug resistant (MDR) isolates, and all *L. monocytogenes* isolates we analyzed were susceptible to 11 different antimicrobials. The frequency of oxacillin (70.8%) and ceftriaxone (62.5%) resistance was relatively high, which might have resulted from the intrinsic resistance to these antimicrobials, consistent with previous studies (6,18,23). In contrast, 20.0% of the *L. monocytogenes* isolates in a Korean slaughterhouse study were MDR (15). In other Korean studies using slaughterhouse, food, and environmental samples, the resistance rates against therapeutic antimicrobials like penicillin, ampicillin, and tetracycline were 53.3–100%, 26.6–97.0%, and 13.3–20.0%, respectively (15,24,25).

Several studies have reported *L. monocytogenes* isolates from foods and environments with resistance to therapeutic antimicrobials (9,13,26–28). Although comparisons are arduous due to differences in the antimicrobial agents and breakpoints used in these studies, *L. monocytogenes* isolates from ruminant listeriosis have relatively lower resistance rates than those reported for food and environmental origins, except for intrinsic antimicrobial resistance. These results might be related to sample source differences and antimicrobial resistance transfer from cross-contamination through the food chain (13,29). The antimicrobial resistance of clindamycin classified into the class of lincosamides, generally used in the treatment of diseases of human and veterinary medicine showed relatively higher intermediate rates (58.3%) than those reported in Poland (34.6%), Italy (29%), and the USA (27%) (12,18,23,26). Furthermore, the resistance of ciprofloxacin-resistant isolates classified into critically important antimicrobial class in WHO also showed slightly higher intermediate rates (29.1%) than those reported in Poland (20.4%) and Italy (24%) (23,26). Considering the acquisition of additional gene mutations, intermediate MIC values in clindamycin and ciprofloxacin widely used in hospitals to treat Gram-positive infections could indicate a possible future shift to resistance phenotypes (26). To our knowledge, there are limited reports of antimicrobial resistance in ruminant clinical listeriosis. Therefore, continuous surveillance of antimicrobial-resistant *L. monocytogenes* is necessary to improve animal and human health. For the treatment of listeriosis, beta-lactams (penicillin and ampicillin), macrolides (erythromycin), aminoglycoside (gentamicin and streptomycin), tetracycline, and trimethoprim/sulfamethoxazole are more effective than ceftriaxone, oxacillin, and clindamycin. Therefore, these agents are recommended as empirical antimicrobials for the treatment of clinical listeriosis.

The significant relevance of antimicrobial resistance patterns among the animal sources, serotypes, PFGE clusters, and STs was not observed in this study, potentially due to the diverse phenotypic and genetic origins of the isolates tested. To elucidate the relevant characteristics that contribute to antimicrobial resistance in *L. monocytogenes*, further studies with larger numbers of isolates from diverse locations and sources are required.

In conclusion, this report is the first comprehensive characterization of *L. monocytogenes* in clinical ruminant listeriosis cases in Korea. Our results indicated that the ruminant *L. monocytogenes* isolates belonged to serotypes 4b, 1/2a, and 1/2b. Based on molecular characterization, clinical ruminant *L. monocytogenes* isolates were genetically diverse and associated with isolates from humans, food, and the environment. According to the clinical signs and histopathological lesions of ruminant *L. monocytogenes* isolates, ruminant listeriosis pathogens, such as ST365, ST9, ST101, ST18, ST224, and ST219, cannot be ignored. Although the ruminant *L. monocytogenes* isolates were susceptible to the primary antimicrobial used for treating human listeriosis, continuous monitoring of the antimicrobial resistance in *L. monocytogenes* isolates is needed. Further research is required to predict the virulence phenotypes and improve the genetic characterization of clinical ruminant *L. monocytogenes* isolates.

## Acknowledgements

We thank all participating investigators and farm owners who provided isolates for Animal and Plant Quarantine Agency.

This work was supported by a grant from the Animal and Plant Quarantine Agency, Ministry of Agriculture, Food and Rural Affairs of the Republic of Korea (Project code no. N-1543069-2015-99-01). The funding body had no role in the design of the study, collection, analysis, and interpretation of data or in writing the manuscript.

## Authors’ contributions

Jongho Kim and Jong Wan Kim: conceptualization, methodology, writing of the original draft, review and editing of the final manuscript.

Ha-Young Kim: conceptualization, supervision, review and editing of the manuscript.

## References

1. Steckler AJ, Cardenas-Alvarez MX, Townsend Ramsett MKT, Dyer N, Bergholz TM. 2018. Genetic characterization of Listeria monocytogenes from ruminant listeriosis from different geographical regions in the US. Vet Microbiol 215:93–97.

2. Wolf-Jäckel GA, Hansen MS, Larsen G, Holm E, Agerholm JS, Jensen TK. 2020. Diagnostic studies of abortion in Danish cattle 2015–2017. Acta Vet Scand 62:1.

3. Papić B, Pate M, Félix B, Kušar D. 2019. Genetic diversity of Listeria monocytogenes strains in ruminant abortion and rhombencephalitis cases in comparison with the natural environment. BMC Microbiol 19:299.

4. Dreyer M, Thomann A, Böttcher S, Frey J, Oevermann A. 2015. Outbreak investigation identifies a single Listeria monocytogenes strain in sheep with different clinical manifestations, soil and water. Vet Microbiol 179:69–75.

5. Dreyer M, Aguilar-Bultet L, Rupp S, Guldimann C, Stephan R, Schock A, Otter A, Schüpbach G, Brisse S, Lecuit M, Frey J, Oevermann A. Listeria monocytogenes sequence type 1 is predominant in ruminant rhombencephalitis. Sci. Rep.:2016 16:(v):36419.

6. Wieczorek K, Osek J. 2017. Prevalence, genetic diversity and antimicrobial resistance of Listeria monocytogenes isolated from fresh and smoked fish in Poland. Food Microbiol 64:164–171.

7. Datta A, Burall L. 2018. Current trends in foodborne human listeriosis. Food Saf (Tokyo) 6:1–6.

8. Du X-j, Zhang X, Wang X-y, Su Y-l, Li P, Wang S. 2017. Isolation and characterization of Listeria monocytogenes in Chinese food obtained from the central area of China. Food Control 74:9–16.

9. Sosnowski M, Lachtara B, Wieczorek K, Osek J. 2019. Antimicrobial resistance and genotypic characteristics of Listeria monocytogenes isolated from food in Poland. Int J Food Microbiol 289:1–6.

10. Wieczorek K, Dmowska K, Osek J. 2012. Prevalence, characterization, and antimicrobial resistance of Listeria monocytogenes isolates from bovine hides and carcasses. Appl Environ Microbiol 78:2043–2045.

11. Wilson A, Gray J, Chandry PS, Fox EM. 2018. Phenotypic and genotypic analysis of antimicrobial resistance among Listeria monocytogenes isolated from Australian food production chains. Genes (Basel) 9:80.

12. Oliveira TS, Varjão LM, da Silva LNN, de Castro Lisboa Pereira R, Hofer E, Vallim DC, de Castro Almeida RC. 2018. Listeria monocytogenes at chicken slaughterhouse: Occurrence, genetic relationship among isolates and evaluation of antimicrobial susceptibility. Food Control 88:131–138.

13. Wang G, Qian W, Zhang X, Wang H, Ye K, Bai Y, Zhou G. 2015. Prevalence, genetic diversity and antimicrobial resistance of Listeria monocytogenes isolated from ready-to-eat meat products in Nanjing, China. Food Control 50:202–208.

14. Abdollahzadeh E, Ojagh SM, Hosseini H, Ghaemi EA, Irajian G, Naghizadeh Heidarlo MN. 2016. Antimicrobial resistance of Listeria monocytogenes isolated from seafood and humans in Iran. Microb Pathog 100:70–74.

15. Oh H, Kim S, Lee S, Lee H, Ha J, Lee J, Choi Y, Choi KH, Yoon Y. 2018. Prevalence, serotype diversity, genotype and antibiotic resistance of Listeria monocytogenes isolated from carcasses and human in Korea. Korean J Food Sci Anim Resour 38:851–865.

16. Thomas EJ, King RK, Burchak J, Gannon VP. 1991. Sensitive and specific detection of Listeria monocytogenes in milk and ground beef with the polymerase chain reaction. Appl Environ Microbiol 57:2576–2580.

17. Kimura M. 1980. A simple method for estimating evolutionary rates of base substitutions through comparative studies of nucleotide sequences. J Mol Evol 16:111–120.

18. Lyon SA, Berrang ME, Fedorka-Cray PJ, Fletcher DL, Meinersmann RJ. 2008. Antimicrobial resistance of Listeria monocytogenes isolated from a poultry further processing plant. Foodborne Pathog Dis 5:253–259.

19. European Committee on Antimicrobial Susceptibility Testing. 2020. Breakpoints tables for interpretation of MICs and zone diameters. Version 10.0, 2020. https://www.eucase.org.

20. Baek SY, Lim SY, Lee DH, Min KH, Kim CM. 2000. Incidence and characterization of Listeria monocytogenes from domestic and imported foods in Korea. J Food Prot 63:186–189.

21. Kathariou S. 2002. Listeria monocytogenes virulence and pathogenicity, a food safety perspective. J Food Prot 65:1811–1829.

22. Swaminathan B, Gerner-Smidt P. 2007. The epidemiology of human listeriosis. Microbes Infect 9:1236–1243.

23. Wieczorek K, Dmowska K, Osek J. 2012. Characterization and antimicrobial resistance of Listeria monocytogenes isolated from retail beef meat in Poland. Foodborne Pathog Dis 9:681–685.

24. Jang YS, Moon JS, Kang HJ, Bae D, Seo KH. 2021. Prevalence, characterization, and antimicrobial susceptibility of Listeria monocytogenes from raw beef and slaughterhouse environments in Korea. Foodborne Pathog Dis 18:419–425.

25. Lee DY, Ha JH, Lee MK, Cho YS. 2017. Antimicrobial susceptibility and serotyping of Listeria monocytogenes isolated from ready-to-eat seafood and food processing environments in Korea. Food Sci Biotechnol 26:287–291.

26. Caruso M, Fraccalvieri R, Pasquali F, Santagada G, Latorre LM, Difato LM, Miccolupo A, Normanno G, Parisi A. 2020. Antimicrobial susceptibility and multilocus sequence typing of Listeria monocytogenes isolated over 11 years from food, humans, and the environment in Italy. Foodborne Pathog Dis 17:284–294.

27. Noll M, Kleta S, Al Dahouk S. 2018. Antibiotic susceptibility of 259 Listeria monocytogenes strains isolated from food, food-processing plants and human samples in Germany. J Infect Public Health 11:572–577.

28. Amajoud N, Leclercq A, Soriano JM, Bracq-Dieye H, El Maadoudi M, Senhaji NS, Kounnoun A, Moura A, Lecuit M, Abrini J. 2018. Prevalence of Listeria spp. and characterization of Listeria monocytogenes isolated from food products in Tetouan, Morocco. Food Control 84:436–441.

29. Wang HH, Manuzon M, Lehman M, Wan K, Luo H, Wittum TE, Yousef A, Bakaletz LO. 2006. Food commensal microbes as a potentially important avenue in transmitting antibiotic resistance genes. FEMS Microbiol Lett 254:226–231.

